# Analysis of the *Streptococcus mutans* proteome during acid and oxidative stress reveals modules of co-expression and an expanded role for the TreR transcriptional regulator

**DOI:** 10.1101/2021.10.25.465825

**Authors:** Elizabeth L. Tinder, Roberta C. Faustoferri, Andrew A. Buckley, Robert G. Quivey, Jonathon L. Baker

## Abstract

*Streptococcus mutans* promotes a tooth-damaging dysbiosis in the oral microbiota because it can form biofilms and survive acid stress better than most of its ecological competitors, which are typically health-associated. Many of these commensals produce hydrogen peroxide, therefore *S. mutans* must manage both oxidative stress and acid stress with coordinated and complex physiological responses. In this study, the proteome of *S. mutans* was examined during regulated growth in acid and oxidative stresses, as well as in deletion mutants with impaired oxidative stress phenotypes, Δ*nox* and Δ*treR.* 607 proteins exhibited significantly different abundance levels across the conditions tested, and correlation network analysis identified modules of co-expressed proteins that were responsive to the deletion of *nox* and/or *treR*, as well as acid and oxidative stress. The data provided evidence explaining the ROS-sensitive and mutacin-deficient phenotypes exhibited by the Δ*treR* strain. SMU.1069-1070, a poorly understood LytTR system, had elevated abundance in the Δ*treR* strain. *S. mutans* LytTR systems regulate mutacin production and competence, which may explain how TreR affects mutacin production. Furthermore, the gene cluster that produces mutanobactin, a lipopeptide important in ROS tolerance, displayed reduced abundance in the Δ*treR* strain. The role of Nox as a keystone in the oxidative stress response was also emphasized. Crucially, this dataset provides oral health researchers with a proteome atlas that will enable a more complete understanding of the *S. mutans* stress responses that are required for pathogenesis, and facilitate the development of new and improved therapeutic approaches for dental caries.

**Importance:** Dental caries is the most common chronic infectious disease worldwide, and disproportionally affects marginalized socioeconomic groups. *Streptococcus mutans* is a considered a primary etiologic agent of caries, with its pathogenicity dependent on coordinated physiologic stress responses that mitigate the damage caused by the oxidative and acid stress common within dental plaque. In this study, the proteome of *S. mutans* was examined during growth in acidic and oxidative stresses, as well in *nox* and *treR* deletion mutants. 607 proteins were differentially expressed across the strains/growth conditions, and modules of co-expressed proteins were identified, which enabled mapping the acid and oxidative stress responses across *S. mutans* metabolism. The presence of TreR was linked to mutacin production via LytTR system signaling and to oxidative stress via mutanobactin production. The data provided by this study will guide future research elucidating *S. mutans* pathogenesis and developing improved preventative and treatment modalities for dental caries.

## Observation

Dental caries remains the most common chronic infectious disease worldwide, and is caused by a dysbiotic dental plaque microbiome that demineralizes tooth enamel via the fermentation of dietary carbohydrates to acid (1). *Streptococcus mutans* is considered a primary etiologic agent of caries due to its exceptional ability to facilitate biofilm formation when provided with sucrose, and its acidophilic niche (2). *S. mutans* employs a robust acid stress response that renders it more acid-tolerant than many of the health-associated commensals that it competes with ecologically. A number of these rival Streptococci produce H_2_O_2_, therefore *S. mutans* must also deal with oxidative stress (3, 4). Numerous studies have examined the role of various genes in these overlapping stress responses and the complex regulatory network that governs them. Previously, our research group identified that the NADH oxidase, Nox, was a linchpin of the *S. mutans* oxidative stress response at the intersection of two oxidative stress regulons (4). Furthermore, the transcriptional regulator of the trehalose utilization operon, TreR, had an unexpected role in oxidative stress and toxin production (5). In this study, mass spectrometry was used to elucidate changes in the *S. mutans* proteome during growth in acid or oxidative stresses, and upon deletion of *nox* or *treR*.

The archetype *S. mutans* strain, UA159 (6), along with the Δ*nox* and Δ*treR* mutant strains were analyzed during tightly-controlled steady-state growth conditions enabled by chemostats set at neutral pH (7), acidic pH (5) and/or sparged with air to maintain an 8.4% dissolved oxygen concentration (i.e. oxidative stress, as described in (4)). Text S1 contains a full description of the materials and methods used in this study. Liquid chromatography-tandem mass spectrometry was performed to examine the proteome of these strains and growth conditions. 1,384 unique proteins were detected across the 8 strains/growth conditions (Table S1). PCA analysis indicated three main clusters of samples: all pH 5 samples, regardless of oxidative stress or genotype; the pH 7 samples without oxidative stress (UA159 and Δ*treR*); and the pH 7 samples under oxidative stress (UA159 + air and Δ*nox*) (Figure 1A). The proteins that were the largest drivers in ordination space towards the pH 5 samples were SpaP, GtfC, GtfD, SMU_63c, while GbpB and AdhE were associated with the pH 7 samples, and Pfl and AtlA were associated with the pH 7 samples under oxidative stress (Figure 1A). Differential expression analyses between pairwise strains/growth conditions is provided in File S1.

**Figure 1:**
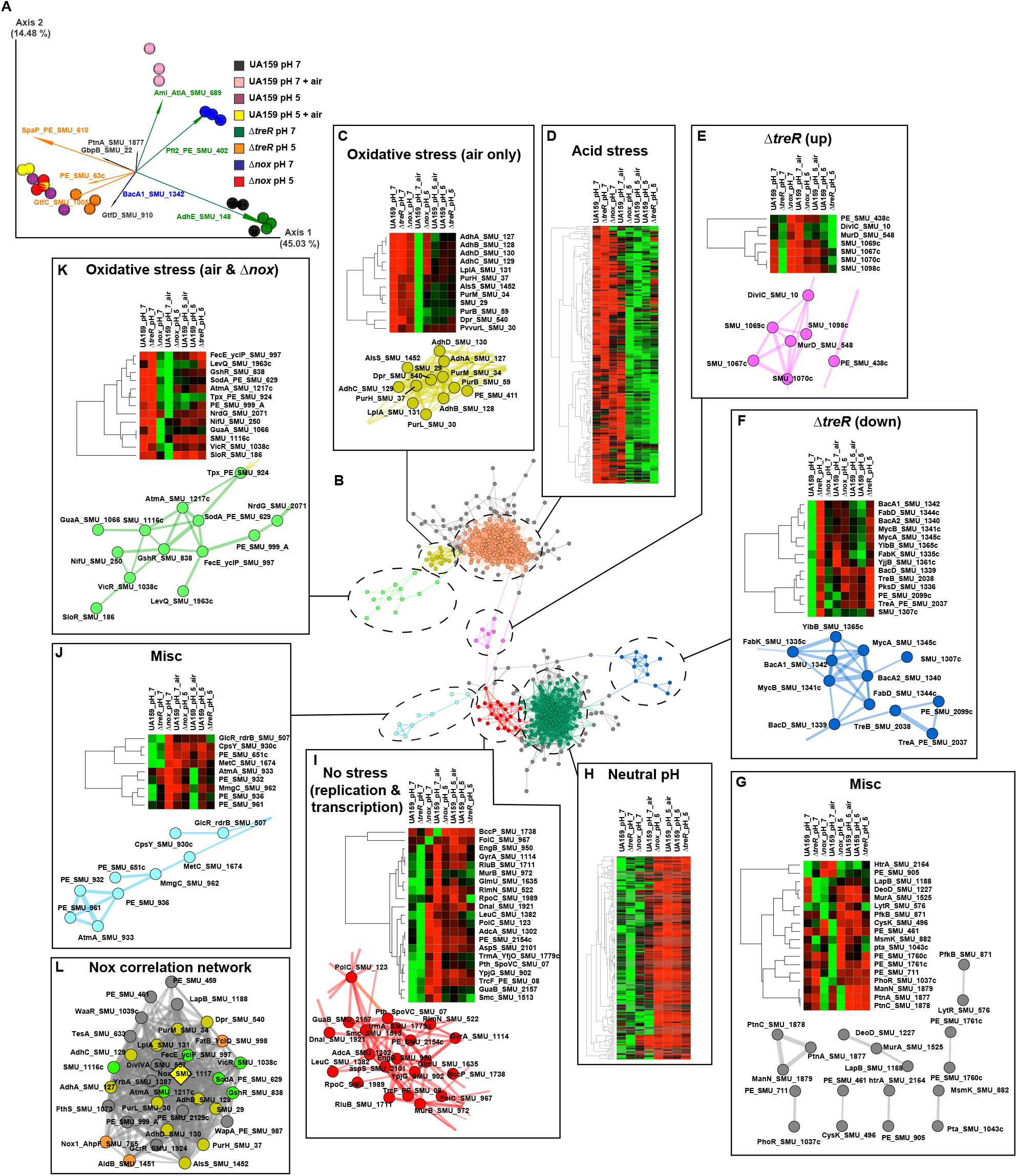
The proteome of *S. mutans* during acid and oxidative stress. **(A)** PCoA biplot of the Bray-Curtis dissimilarity between samples of the indicated strain and growth condition, represented by the colored spheres. Feature loadings (i.e. proteins driving distances in ordination space) are illustrated by the vectors, which are labeled with the cognate feature name and colored based on that feature’s cluster in Panel B. **(B)** Clustering of *S. mutans* proteins into co-expression clusters. Protein association network illustrating co-expressed proteins. Prior to clustering, proteins were filtered for significant differences using an uncorrected ANOVA p < 0.01 (607 proteins). Correlations (edges) with a Spearman’s ρ > 0.8 are shown and only positive correlations were considered. Edge width is representative of Spearman’s ρ. Clusters were manually selected as indicated by the node color. **(C-K)** Sub-clusters are annotated with a heatmap indicating protein abundance across the 8 strains/growth conditions. Heatmap rows are clustered by Spearman’s ρ. A version of the full network with each node labeled is available in Figure S1, and versions of the Acid Stress and Neutral pH heatmaps with each row labeled are available in Figure S2. A pairwise correlation table of all proteins is provided in Table S2. A heatmap illustrating expression of the 54 proteins that were differentially expressed based on ANOVA, but did not have significant correlations with other proteins, is provided in Figure S3. **(L)** Proteins that correlate with Nox when the Δ*nox* samples are not included in the network analyses. The Δ*nox* strain data likely obscured proteins that correlate with Nox, therefore the correlation network analysis was repeated without the Δ*nox* data. The network shown here is a sub-cluster of all 33 proteins significantly correlating with Nox expression. Nox is represented by the yellow diamond, all other nodes are colored by the sub-cluster determined in Figure 1B-J. Edge is representative of Spearman’s ρ. Only positive correlations with ρ > 0.8 are shown.

Correlation network analysis was performed to observe modules of co-expressed proteins under the various conditions (Figure 1B). This analysis revealed two large clusters of proteins associated with elevated abundance at either pH 5 or pH 7 (Figure 1D and H). Several smaller sub-clusters were associated with other discrete expression profiles such as oxidative stress or deletion of the TreR regulator (Figure 1CEFGIJK). A cluster associated with oxidative stress, either through addition of air or deletion of *nox*, included many of the well-established proteins of the oxidative stress tolerance response, including Tpx, GshR, Sod, SloR, and VicR (Figure 1K). An adjacent cluster of proteins, including the Adh operon, as well as Dpr, AlsS, and much of the purine biosynthesis gene cluster, had elevated abundance at pH 7 with air, but not in Δ*nox* (Figure 1C).

Intriguingly, two sub-clusters displayed expression profiles specifically affected by the presence of the TreR regulator. DivIC and MurD, involved in cell wall synthesis and cell division, as well as the autoregulatory LytTR system, SMU.1069-1070, had increased expression in the Δ*treR* strain (Figure 1E). SMU.1069-1070 exhibits crosstalk with the more well-characterized LytTR systems, HdrRM and BrsRM, known to regulate competence and bacteriocin production (7, 8). Since TreR and trehalose operon expression play a role in competence (9), and the production of mutacins IV, V, and VI (5), through unknown mechanisms, signaling through SMU.1069-1070 is an attractive hypothesis. Although the mutacin IV, V, and VI NRPS products themselves are too small to be detected by the proteomics analysis employed here, further evidence linking TreR to mutacin production was observed. Several proteins within mutacin biosynthetic gene clusters (BGCs) did have significantly decreased abundance in the Δ*treR* strain, including CopYAZ (mutacin VI BGC) and SMU.1904 and SMU.1910 (mutacin V/CipB BGC) (File S1).

Meanwhile, the proteins from the trehalose operon itself, as well as the large mutanobactin BGC (SMU.1334-1349) were reduced in the Δ*treR* strain (Figure 1F). This further confirmed that in *S. mutans*, TreR serves as an activator of tre operon expression, rather than as a repressor, as seen in other species (5). Mutanobactin, a non-ribosomal lipopeptide, appears to have a role in helping *S. mutans* deal with oxidative stress (10). Therefore, it is possible that reduced abundance of the mutanobactin BGC may explain the impaired ROS tolerance in the Δ*treR* strain. Interestingly, Nox and TreR did not appear in the correlation network, likely due to their absence in deletion mutant strains obscuring correlations. In repeated correlation analysis with the deletion mutant samples removed, Nox expression was tightly-correlated with 33 co-expressed proteins, mainly from the clusters of genes associated with oxidative stress, further confirming its role as a keystone protein in the *S. mutans* oxidative stress response (Figure 2L). Contrarily, in the reanalysis, TreR only had one protein correlation with ρ ≥ 0.8, SMU_690. Since TreR did not exhibit strong correlation with other proteins, but its absence had a major effect on the abundance of a number of proteins, it seems modulation of transcriptional regulatory activity for TreR, rather than just TreR expression level, is likely to be key in its role as a regulator.

**Figure 2:**
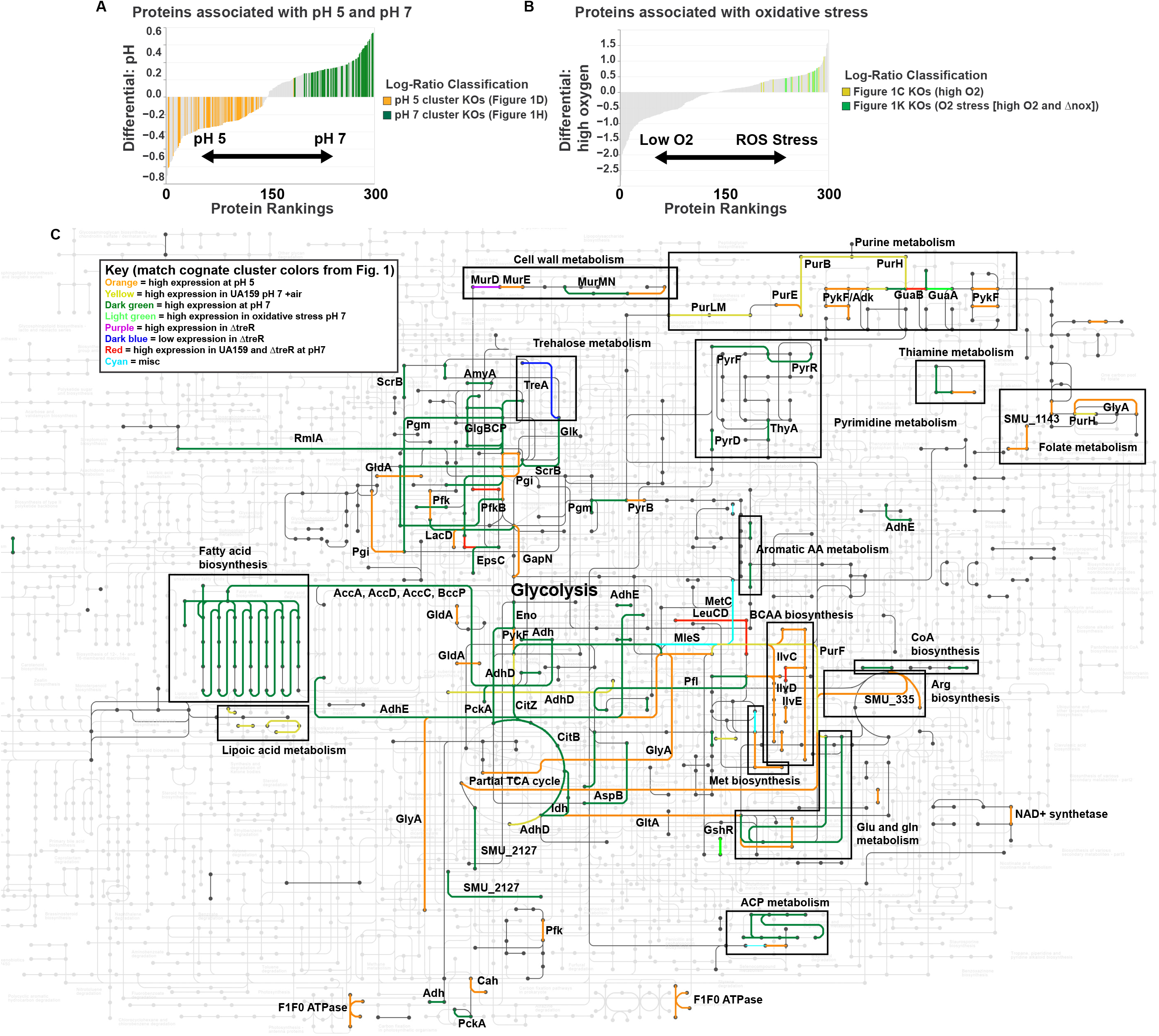
Metabolic modules of the *S. mutans* acid and oxidative stress responses. **(A)** Differential ranking of proteins associated with pH 5 vs pH 7. Songbird was used to rank proteins with respect to pH, and Qurro (13) was used to visualize the resulting ranks (only the top and bottom 150 proteins are shown). Proteins with known KOs in the sub-clusters shown in Figures 1D and 1H are highlighted in orange and dark green, respectively. **(B)** Differential ranking of proteins associated with high O_2_ (UA159 + air and Δ*nox*) vs. low O_2_ (UA159 and Δ*treR*). Songbird was used to rank proteins with respect to high vs low O_2_, and Qurro (13) was used to visualize the resulting ranks (only the top and bottom 150 proteins are shown). Proteins with known KOs in the sub-clusters shown in Figures 1C and 1K are highlighted in yellow and light green, respectively. **(C)** Metabolism of *S. mutans* during acid and oxidative stress. All proteins from the sub-clusters shown in Figure 1C-K with known KOs were overlaid on to a map of the known metabolism of *S. mutans* using KEGG Mapper (https://www.genome.jp/kegg). Colors of each sub-cluster from Figure 1 are maintained, as described in the Key.

Differential rankings (11) were utilized to determine the proteins most associated with acid and oxidative stress. KOs from the sub-clusters associated with pH 5 and pH 7 (Figures 1D and 1H) made up the majority of the proteins associated with the cognate pH (Figure 2A), while proteins from the sub-clusters associated with oxidative stress (Figures 1C and 1K) were in fact correlated with the associated growth condition, based on supervised methods (Figure 2B). To further examine the impact of the genotypes and growth conditions on *S. mutans* metabolism, proteins with associated KO numbers from the sub-clusters in Figure 1C-J were overlaid onto a map of the metabolism of *S. mutans* UA159 using KEGG Mapper (https://www.genome.jp/kegg/) (Figure 2). Table S3 provides a table of KO numbers and colors to be used by the reader to generate an interactive version of the metabolic map shown in Figure 2C using KEGG Mapper Color. Many of the large-scale trends observed were in-line with previous, transcriptomic and proteomic observations (3, 12). These included increased abundance of proteins involved in fatty acid biosynthesis, the partial TCA cycle and pyrimidine metabolism at pH 7, and increased abundance of proteins involved in arginine deiminase, BCAA biosynthesis, purine metabolism, and the F_1_F_0_ ATPase at pH 5. Overall, this updated perspective of the *S. mutans* proteome provides a comprehensive interpretation of how this organism deals with acid and oxidative stress, permitting its key role in the dysbiosis that leads to caries pathogenesis. This study also highlights several principal avenues for future research, including the importance of the TreR regulator.

## Data Availability

The raw mass spectrometry output files are available in the MassIVE Repository (massive.ucsd.edu) with the accession number MSV000088252.

## Supplemental Figure and File Legends

**Figure S1: Correlation network of the *S. mutans* proteome.** The complete Figure 1B network, with each node is labeled with the cognate protein name. Clustering of *S. mutans* proteins into co-expression clusters. Protein association network illustrating co-expressed proteins. Prior to clustering, proteins were filtered for significant differences using an uncorrected ANOVA p < 0.01 (607 proteins). Correlations (edges) with a Spearman’s ρ > 0.8 are shown and only positive correlations were considered. Edge width is representative of Spearman’s ρ. Clusters were manually selected as indicated by the node color.

**Figure S2: Expression profiles of proteins associated with pH 5 (A) or pH 7 (B).** These are expanded versions of the heatmaps appearing in Figure 1 panels D and H, with each row labeled. Rows are clustered by Spearman’s ρ.

**Figure S3: Expression profile of differentially-expressed genes that had no significant correlations.** Heatmap showing the expression of proteins that made the differential expression ANOVA p ≥ 0.5 cutoff, but did not have any correlations with other proteins with a Spearman’s ρ ≥ 0.8. Rows are clustered by Spearman’s ρ.

**Table S1: Normalized abundances of detected proteins**

**Table S2: Spearman’s Rank correlations between differentially-expressed proteins.**

**Table S3: KO and color list to generate interactive *S. mutans* metabolism map using KEGG Mapper – Color** (https://www.genome.jp/kegg/mapper/color.html)

**File S1: Excel file containing pairwise log_2_ fold-changes and p-values for each protein between all 8 strain/growth conditions**

**Text S1: Supplemental Materials and Methods**

## Acknowledgements

The authors thank Kevin Welle and the Mass Spectrometry Resource Lab at the University of Rochester Medical Center for performing the proteomics analysis. This study was supported by NIH/NIDCR grants R01-DE013683 (R.G.Q.) and K99-DE029228 (J.L.B.).

